# Two complementary circuits for spatial navigation in mouse: evidence from Fos mapping

**DOI:** 10.1101/2023.10.13.562274

**Authors:** Edyta Balcerek, Urszula Włodkowska, Rafał Czajkowski

## Abstract

We designed and implemented a novel spatial behavioral protocol based on the figure 8 maze paradigm. This solution employs a fully automated closed-loop system using visual contexts for guiding behavior. It allows high throughput testing of large cohorts of animals. This automated approach highly reduces the effect of stress, but also eliminates the influence of rule learning and the involvement of working memory. Here, we show that training mice in the apparatus leads to change in strategy from internal tendency to alternate into navigation based exclusively on visual cues. This effect could be achieved using two different protocols: prolonged alternation training, or a flexible protocol with unpredictable turn succession. We provide evidence of opposing levels of engagement of hippocampus and retrosplenial cortex after training mice in these two different regimens. This supports the hypothesis of the existence of parallel circuits supporting navigation, one based on the well described hippocampal representation, and another, RSC-dependent.

## Introduction

Choosing the optimal strategy for wayfinding and spatial exploration is one of the fundamental tasks for all animals. Maintaining the correct balance between exploiting the already known territories and exploring the new ones is a continuous process, supported by a dedicated network of brain structures. Historically, the research on forming, updating and maintaining the spatial map has been centered around the hippocampal formation. More recent studies suggested that other structures might also participate in this process, with various degrees of independence from the hippocampus (HPC). One of the heavily investigated but still underappreciated regions is the retrosplenial cortex (RSC) [1]. It connects extensively with the hippocampal formation, but also independently with other areas, including the anterior cingulate, visual and parietal cortices [2]. We have previously contributed to mapping the relevant connectivity of the RSC and revealed a direct functional input from this structure into the deep layers of medial entorhinal cortex (MEC) [3, 4]. These layers serve as the output of the MEC-HPC loop, therefore any information directly modulating their activity might affect the outcome of hippocampal processing. We recently confirmed that the HPC and RSC input converge on the same population of MEC neurons [5], suggesting the existence of two parallel circuits sharing the same output [6]. Others have shown that the hippocampus might exert an inhibitory effect on the RSC activity via direct inhibitory projections from CA1 and subiculum, preventing competition between the two structures [7]. We have also shown that a measurable behavioral output in a spatial exploration paradigm could be produced by a range of alternative brain networks. In a seemingly homogeneous population of animals, different spatial strategies might spontaneously emerge and develop [8]. In this paper we investigated the possibility of engaging a specific spatial memory circuit as a result of the training protocol used. We designed and implemented an automated, semi-autonomous system for spatial memory tests. It is based on the T-Maze paradigm [9–12], with modifications allowing for reliable dependence on visual cues. It allows for a relatively high throughput with minimum human intervention. Using two different regimens, we trained two groups of animals to perform the task at comparable levels. We then mapped the expression of the immediate early gene Fos in the hippocampus and the retrosplenial cortex.

## Materials and methods

### Animals

Fourteen young (12 weeks old) male, wild-type C57BL/6 mice were used for this experiment. One animal was excluded during training due to health issues. Experiments were performed with individually-housed mice. Animals were kept on a reversed 12-h light/ dark cycle, with the dark phase starting at 7 am. All behavioral procedures were conducted during the dark phase. Mice had ad libitum access to food and water before the experiment. Along with the beginning of experimental procedures, minimal food restriction (3,5g daily/ mouse) was implemented in order to raise animals’ motivation for the food reward. Average daily body weight was monitored for each animal and never dropped below 90% of initial body weight. All experiments were approved by the 1st Local Ethical Committee in Warsaw (code: 1397/2022). The experimental protocol followed the European Communities Council Directive and The Law on the Protection of Animals Used for Scientific or Educational Purposes.

### Apparatus

A fully automated figure-8 maze apparatus, (external dimensions of the enclosure: 80 x 45 cm, wall height 25 cm) was designed and in-house manufactured (Fig. 1A). All front walls and the stem walls were made of transparent plexiglass, all other walls and the floor were opaque. The corridor width was 10 cm. The length of the stem section was 25 cm, the length of each arm was 35 cm. Seven pneumatically operated doors were installed in order to enable continuous movement of the animals and to force a desired turn direction when necessary (Fig. 1B). Visual information was presented using an array of triangular LED panels (Aurora system, Nanoleaf). Two different visual contexts (Fig. 1C) were displayed in front of the maze. Both contexts varied in terms of colors and spatial arrangement of LED panels. Each context was associated with a specific turn direction. Sweet condensed milk solution (approx. 10 *μl*) was used as the food reward. Reward was administered at the front corners from dispensers with electromagnetic valves via 18G blunt needles.

**Fig 1.**
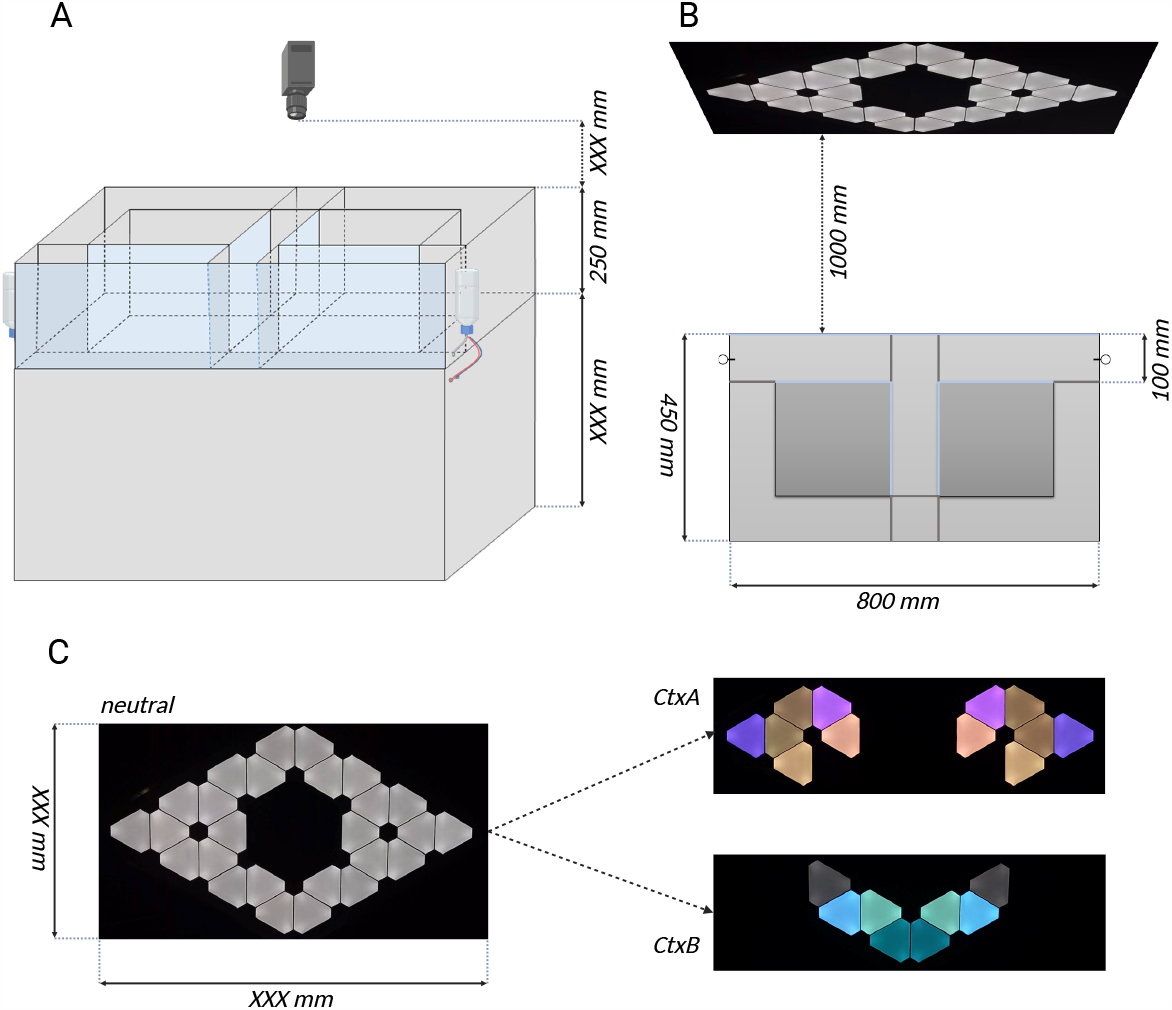
Figure-8 maze apparatus (see text for details). A) Schematic front view with camera and reward dispensers locations. B) Schematic top view with LED panels placed 1000mm ahead of the transparent, front wall showing natural context used for habituation protocol as well as for starting and ending of each daily session of behavioral procedure. Transparent walls are light blue colored. All inner doors are pneumatically movable in order to enable continuous movement of the animals and to force a desired turn direction when necessary. C) Realistic view of displayed visual cues. Neutral context (left) showing the full array of triangular LED panels, CtxA (right, top) associated with right turn, CtxB (right, down), associated with left turn.

### Habituation

For all behavioral procedures mice were handled for at least 14 days prior to the onset of training. During the last 7 days, handling sessions were designed to mimic the final experimental protocol, including appropriate light and sound changes, conditions of the experimental room and getting used to the apparatus. In order to completely eliminate the novelty stress associated with the reward, mice were exposed to sweet reward (condensed milk) in the home cage for 4 min/day on four consecutive days. Milk was applied through the same dispensers as used in the apparatus during training. Each time the mouse approached the dispenser, it received a drop of milk (approximately 10 *μl*). When all animals developed a stable approach behavior towards the reward immediately after inserting the dispenser, the activity was completed. Few hours after milk stimulus exposure mice were habituated to the figure-8 maze apparatus, where subsequent stimuli were introduced gradually. For two consecutive days animals were able to move freely inside the maze for 4 minutes/day with no door movement and the reward was automatically provided each time the mouse approached the dispenser area. During the next two days mice were placed in the apparatus for 4 minutes/day with some (5 of 7) motorized doors activated and the reward given only once per specific arm entry. Each exposure to the maze apparatus prior to the onset of training was accompanied by the neutral context displayed on LED panels, as the only light source (Fig. 1C).

### Experimental Protocol

After the habituation period mice underwent 12 days of spaced training (Fig. 2A-C). Each trial consisted of two phases with the mouse passing through the maze in a continuous manner. Each trial began with the forced phase, with the animal starting in the center zone, entering the single available arm of the maze and immediately after collecting the reward returning to the center zone. Then the second phase of the trial (choice phase) started with both arms open so that the animal could make a direct choice and decide which arm to enter. Both runs were rewarded, but milk was given only in the correct arm, according to the visual context displayed in front of the maze. Contexts were defined specifically for left and right turns respectively (Fig. 1C) and displayed during both runs (forced and choice) giving the opportunity to navigate based on external directions at each run. Each animal underwent a progressively increasing number of forced-choice trials per day. Training started with 8 trials per day and reached the full number of trials - 25 on the 10th day of training. Two groups of animals (“Alternation” and “Modified”) were trained in two different navigational regimes. Alternation training was based on a repetitive scheme of contexts displayed in alternating order. This strategy takes advantage of the natural tendency of rodents to explore unvisited territory [13]. Each forced run was immediately followed by the choice run with the changing of visual context. The correct response for the animal was to choose the opposite arm from the one it visited on the previous (forced) run (win-shift strategy) (Fig. 2B). Modified training required flexible response to visual stimuli. In half of the trials, the schematic, alternating sequence described above was replaced with trials in which the system displays identical context in both forced and subsequent choice runs. The rewarded response is for the animal to choose the same arm it had visited on the previous (forced) run. Such complex trials (with the same context displayed in both forced and choice runs) were programmed among alternating trials with a frequency of 50% of all trials (Fig. 2C). The experimental design is summarized in Fig. 2A-C. We designed a closed-loop system in which the position of the animal was tracked. After the animal made a choice, the doors to both arms were closed and the reward was dispensed only when the chosen arm was the correct one. The final number of right and left turns (corresponding with visual cue) was equal each day. Both training groups were tested on the 13th day of the experiment in an identical manner. The test was very similar to the strategy of “Modified” training with one exception: all 25 forced-choice trials of flexible response to changing visual cues were rewarded to avoid extinction.

**Fig 2.**
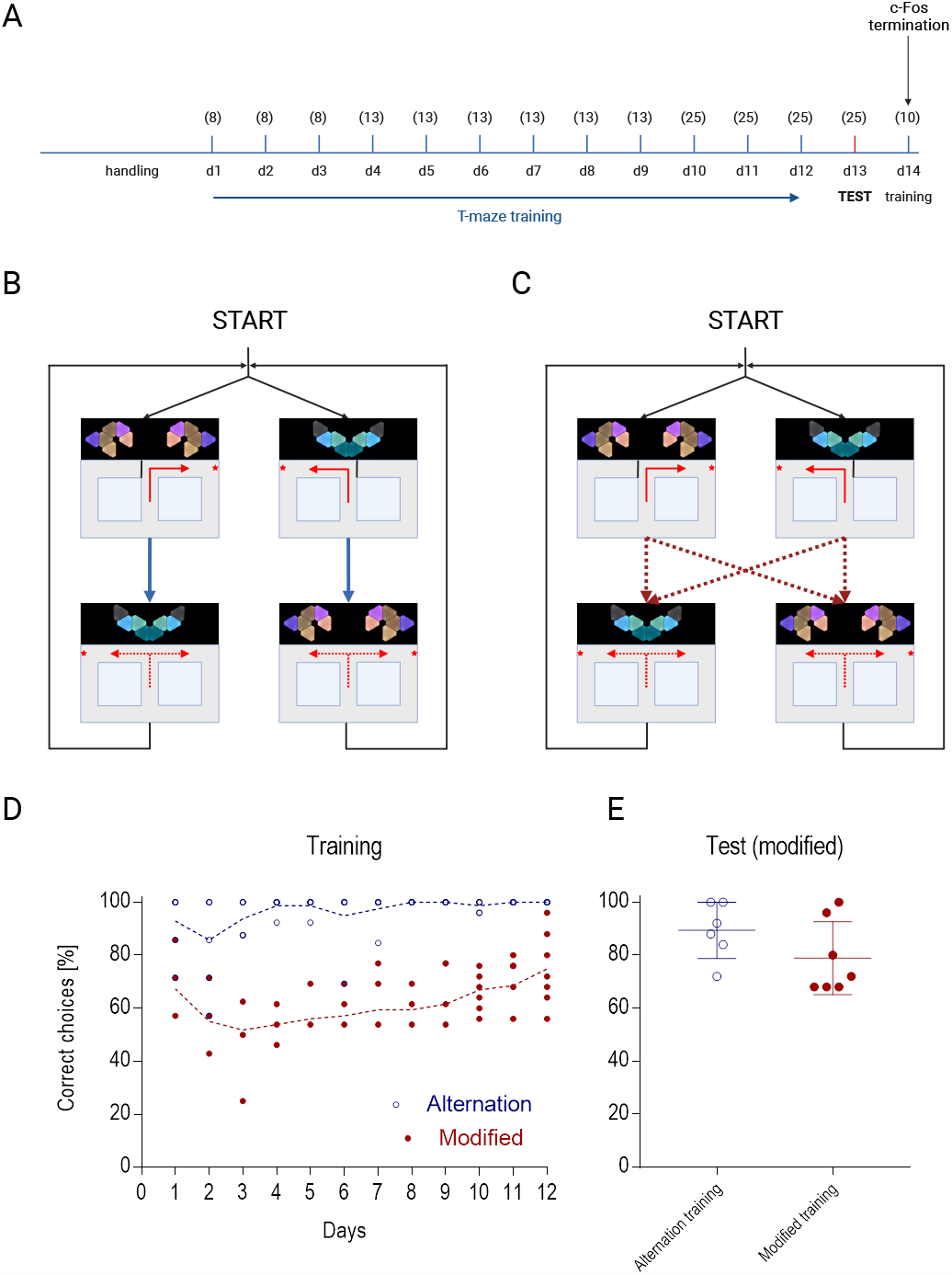
Behavioral experiment. A) Timeline representation of the experimental design (see Methods for details). Numbers in brackets indicate subsequent numbers of forced-choice trials at each day of spaced training. Final number right and left turns (corresponding with visual cue) was equal on each day. B) Schematic representation of “Alternation” training strategy. A rigid order of turns was imposed. C) Schematic representation of “Modified” training strategy. Any sequence of turn directions was possible between forced and choice phase, while maintaining an equal proportion of every possible sequence. D) Learning curve for “Alternation” and “Modified” groups. Data points show the percentage of rewarded decisions (congruent with the displayed context) by individual animals in each session. E) Test session in “Modified” layout. Data points show the percentage of decisions congruent with displayed context.

### Fos induction

In order to induce the expression of Fos protein, 24 hours after the behavioral test, each animal was subjected to 10 trials (20 subsequent runs) in the “Modified” layout (Fig. 2A). Ninety minutes after the beginning of the task, each animal was quickly sedated with a mixture of isoflurane (5%) in conjunction with pure oxygen, followed by intraperitoneal administration (30G needle) of ketamine (75 mg/kg) and medetomidine (0.5-1 mg/kg) mixture. Then mice received an overdose of sodium pentobarbital (150 mg/kg) and were perfused transcardially with phosphate-buffered saline (0,1M PBS, 50ml, RT) with heparin (10 000 units/l) followed by paraformaldehyde (PFA) perfusion (4% paraformaldehyde solution in 0.1 M PBS, 150 ml, 4°C). The brains were collected and stored for 24 hours in PFA, followed by sucrose solution (30% sucrose in 0.1M PBS). Brain slices were cut coronally (40 *μm*) in a cryostat and subjected to histological analysis.

### Immunostaining

In order to detect Fos protein, free-floating coronal brain sections were first washed 3 x 10 min in 0.1M PBS (40 rpm). Background staining (activity of aldehyde group) was diminished by 15 min incubation in 1% NaBH4 in 0.1M PBS (2 rpm) followed by subsequent washes in 0.1M PBS (1 x 10 min, 3 x 5 min, 30 rpm). Endogenous peroxidase activity was blocked by incubating the brain sections in solution of 3% hydrogen peroxide and 70% methanol in distilled water for 10 min (10 rpm), followed by four subsequent washes in 0.1M PBS, 5 min each (30 rpm). Sections were blocked in 5% normal goat serum (NGS) in 0.3% PBST (phosphate-buffered saline solution with a 0,3% concentration of triton x-100) for 4h and incubated 48h at 4°C with solution of primary c-Fos rabbit polyclonal antibody diluted 1:1000 (cat. nr: SYSY 226003) in 0.3% PBST with 5% NGS. Incubation was stopped with 0,3% PBST / 5% NGS (5 min, 30 rpm), sections were washed three times in solution of 0,3% PBST / 2% NGS, 10 min each (30 rpm). Then a secondary biotinylated goat anti-rabbit antibody was applied in PBST / 2% NGS (1:200; Vectastain, Vector Laboratories, Burlingame, CA, USA) (75 min, RT, 10 rpm). The sections were then washed (5 x 5 min, 20 rpm) and processed with avidin-biotinylated horseradish peroxidase complex (Vectastain Elite Kit, Vector Laboratories) in 0.3% PBST for 1 h at room temperature (2 rpm), followed by subsequent washes in 0.3% PBST (4 x 5 min, 15 rpm). The reaction was visualized using diaminobenzidine (DAB, Vector Laboratories, Burlingame, CA, USA), enhanced by nickel salt and was stopped by PBS washes (4x). Sections were mounted on glass slides and left overnight. Sections were then further dehydrated in graded alcohol concentrations (ethanol), defatted by xylene and coverslipped with water-free mounting medium (Entellan).

### Fos cells identification

Digital images were captured with an Olympus VS110 microscope. High-resolution scans of the whole brain slice specimen were taken. To acquire the best precision in selecting the region of interest digital histological sections were overlaid with relevant transparent atlas figures shaping the frame of structures borders. Frames of brain coronal borders were created in the GNU Image Manipulation Program (Free Software Foundation). Image analysis was performed with the ImageJ platform followed by custom-made semi-automated scripts detecting locations of c-Fos positive cells. c-Fos expression was reflected by the density of c-Fos immunopositive nuclei that were labeled above a set threshold. The threshold was determined individually for each region of interest: rRSA, cRSA, rRSG, cRSG, dCA1, dCA3, dDG and was kept the same among all mouse samples. Brain sections were identified according to the atlas of Paxinos and Franklin (14). For each brain, counts were taken from seven anterior-posterior levels centered around the target AP coordinate (rostral part of RSC (rRSA, rRSG) -1.50 mm; caudal part of RSC (cRSA, cRSG) -2.90 mm; dorsal HPC (dCA1, dCA3, dDG) -1.50 mm from bregma respectively). The total number of c-Fos-immunopositive nuclei from both right and left sides were considered. To calculate the density [cells/mm2] of c-Fos positive cells in each area, the number of c-Fos stained cells in a given field was counted and divided by the area that was occupied by the given field.

### Data analysis

All results were analyzed by researchers unaware of exact experimental conditions. Datasets were aggregated in Excel and further analyzed in Prism v7.01 (GraphPad Software). To compare learning progress in the Alternation and Modified groups, Two way repeated measures ANOVA was applied, with the post-hoc Tukey test to analyze individual time points. To compare the test performance, t test was applied. For c-Fos density comparison Holm-Sidak multiple t test was used. All values are expressed as means with standard deviation indicated.

## Results

We first verified the ability of mice to perform the visually guided version of the figure-8 maze task in the automated apparatus following two different training protocols. In both versions of the task mice were expected to associate the displayed spatial context with the rewarded turn direction (see Methods for details). In the “Alternation” training scheme, apart from the spatial guidance, the win-shift strategy was imposed on the animals, with the reward location changing between forced and choice phase for each trial. Accordingly, context was switched between the forced and choice phase, always remaining congruent with the turn direction. In this case, contextual information was additionally supported by the rule learning. As expected, mice (n=6) acquired the task within no more than three sessions and remained at the asymptotic level for the rest of the training (accuracy in the last training session: 100%, Fig. 2D). In the “Modified” version, with no predictable alternation, the win-shift strategy was correct only in 50% of the trials. In the remaining half, the win-stay strategy was imposed and the animals had to reenter the previously visited arm in order to collect the reward. For each phase of each trial the goal location was aligned with specific visual context, therefore the ability to flexibly suppress the win-shift strategy and to rely solely on the contextual information was necessary for successful learning. Under such a regimen, mice (n=7) initially performed around chance level. The apparent lack of consistent reinforcement led to an initial decrease of performance, reaching a minimum at Session 3 (average accuracy: 51.8%, s.d.=12.37). However, over the course of Sessions 4-12 the group gradually learned (p=0.0002, Tukey’s test). At the conclusion of the training phase some animals in this cohort improved to almost match the performance of the “Alternation” group (average accuracy in the last training session: 74.9%, s.d.=12.95, Fig. 2D).

We then tested the ability of both training groups to solve the task only in the “Modified” layout, with 50% of the trials requiring reentry to the previously visited arm. During the test both arms were rewarded in the choice trial to avoid memory extinction. As expected, in the “Modified” group we observed performance similar to the last training session (average accuracy: 78.8%, s.d.=13.8, Fig. 2E). Unexpectedly, the “Alternation” group, despite following only the schematic, win-shift protocol during training, also performed at a high level in the “Modified” test (average accuracy: 89.3%, s.d.=10.6, Fig. 2E) and no statistical difference between groups was observed (p=0.159, t test).

After 24 hours we performed another training session (20 trials) in the “Modified” layout in order to induce c-Fos expression in the relevant brain areas responsible for spatial learning. 90 minutes after beginning of the task, mice were perfused and brains were processed for Fos immunohistochemistry. We focused the analysis on two relevant regions, previously identified as involved in spatial memory, navigation and wayfinding (Fig. 3A). We analyzed the dorsal hippocampus (dentate gyrus, CA1 and CA3 fields) and the retrosplenial cortex (granular and agranular) at two coronal planes (rostral and caudal, see Methods for details). We compared expression levels between mice trained in two learning protocols, but tested in the same conditions. Our analysis revealed that the dorsal hippocampus becomes highly activated in the group that was trained in the “Modified” protocol (Fig. 3B). Density of c-Fos immunopositive cells was much higher in the dorsal part of the CA1 field (2456 cells/mm2, s.d.=1178) compared to mice from the “Alternation” group (848 cells/mm2, s.d.=428, adjusted p=0.027, Holm-Sidak multiple t test). A similar trend was observed in the CA3 region, but when adjusted for multiple comparisons, the difference did not reach statistical significance (1765 cells/mm2, s.d.=794 vs. 934.7 cells/mm2, s.d.=457, adjusted p=0.088). The dentate gyrus, a subregion of the hippocampal formation usually showing sparse Fos expression, does not seem to be more active in the alternation group (549 cells/mm2, s.d.=484) compared to the modified protocol (836 cells/mm2, s.d.=392, adjusted p=0.262).

**Fig 3.**
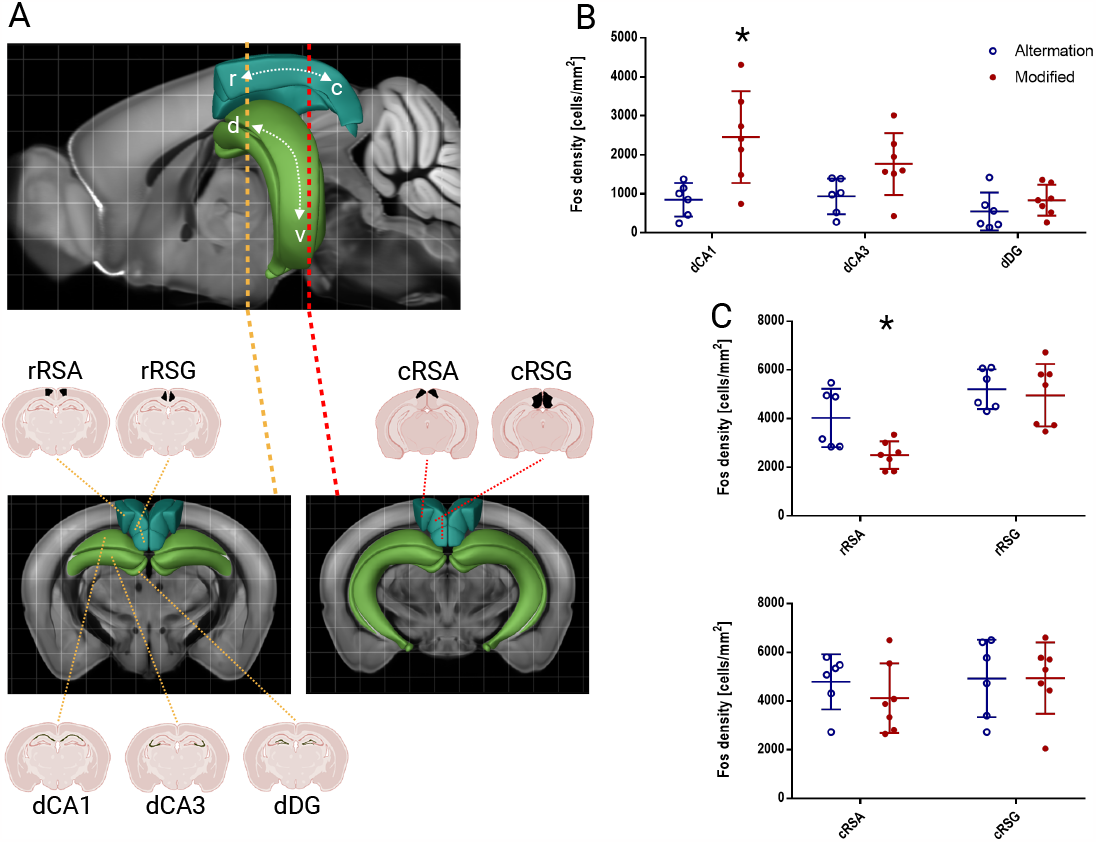
Comparison of c-Fos-positive cell density between training protocols across brain regions. A) Visualization of the position of central slices for each region of interest in the mouse brain using Allen Brain Atlas. In the upper panel, hippocampus and retrosplenial cortex are marked with colors, and the rostro-caudal position of the slices is indicated. The lower panel shows the position of ROIs for c-Fos positive cell density collection on the coronal slices. B) Density of c-Fos immunopositive cells in the ROIs of dorsal hippocampus, compared between both experimental groups. A statistically significant difference between training protocols has been revealed for CA1, where c-Fos expression proved to be higher in mice trained in the “Modified” protocol (p=0.027, Holm-Sidak multiple t test). A similar trend (however, with no statistical significance, p=0.088) was observed for CA3. C) Group differences of activation of the rostral part of granular (rRSG) and agranular (rRSA) subdivisions of RSC. A significant suppression of the c-Fos expression in the “Modified” protocol was observed for rRSA (p=0.023, Holm-Sidak multiple t test) Similar analysis for the caudal part or RSC does not show a visible difference. This result proves the existence of activation gradient along the rostro-caudal axis for agranular retrosplenial cortex.

Similar analysis performed for the retrosplenial cortex included two coronal planes (see Methods), since functional gradients have been previously reported for this structure. In the “Alternation” group we observed an almost uniformly high activation across RSC subregions (Fig. 3C), with no differences along the rostro-caudal axis (rRSA, 4031 cells/mm2, s.d.=1200, rRSG, 5209 cells/mm2, s.d.=814, cRSA, 4789 cells/mm2, s.d.=1129, cRSG, 4925 cells/mm2, s.d.=1590). In the “Modified” group, a the activity of rostral RSA was markedly suppressed (2494 cells/mm2, s.d.=567, inp=0.023), while RSG and caudal part of RSA were activated at a similar level as the “Alternation” group (rRSG, 4958 cells/mm2, s.d.=1284, cRSA, 4114 cells/mm2, s.d.=1426, cRSG, 4938 cells/mm2, s.d.=1464).

We also verified the correlation of behavioral response with Fos levels in all brain regions of interest. Spatial learning ability for individual mice defined as task acquisition (measured by percent of correct turns according to the visual cues in test) was not corresponding with the c-Fos level in the “Modified” group. On the other hand, the performance of the “Alternation” group during the test trial was strongly correlated with the Fos expression level in rostral RSG (Spearman correlation, r=0.899, adjusted p=0.028).

## Discussion

Studies of spatial learning and navigation based on the allothetic information have always been considered a technical and conceptual challenge when applied in the mouse model. In this work, we successfully implemented a novel behavioral task that tests the ability of mice to follow the correct path and to retrieve a reward, based solely on the spatial context presented. Although similar automated maze devices were constructed previously [11, 12, 15–18], and the T-Maze protocol has been tested in a plethora of previous studies [10, 19–23], it was always used in the win-shift arrangement, where spatial information merely supported the rule learning. The animals had to remember the given spatial orientation and utilize it to choose the opposite arm during the choice phase. Notably, in such cases, working memory also played an important role in solving the task. We show it is possible to dissociate the spatial memory from the rule learning and to completely eliminate the working memory component, either during the training phase, or selectively in the behavioral test. To the best of our knowledge, only one similar attempt has been successfully shown [24], although the VR-based model used in this study in combination with an immobilized mouse eliminated active movement and limited the range of practical applications. In our approach, high throughput experiments are possible, with the capability of training and testing up to tens of animals in one day session. This opens a possibility to use it as a broad screen for the mechanisms of spatial memory impairments in the mouse model, similarly to the solutions reported before [12].

Considering the fact that our behavioral test session revealed two populations with matching behavioral performance, we decided to compare the relative involvement of two key brain regions responsible for spatial memory and navigation: dorsal hippocampus and retrosplenial cortex in task completion. We used the expression patterns of c-Fos immediate early gene as a proxy for behaviorally relevant neuronal activation [6]. Two distinct behavioral approaches used for training allowed for within-experiment comparisons, without the need for additional naive control. Interestingly, there was a marked difference in the hippocampal activity in the group that followed the flexible learning and working memory-independent regimen already during training (“Modified” group). It was particularly prominent in the CA1 region, although a similar trend could be observed along the entire trisynaptic pathway.

The limited scale of performed analysis does not allow for definite answers, but it could be speculated that the hippocampal formation is activated when spatial navigation in the known environment is driven by the need for flexible location decisions, determined by competing or overlapping extramaze cues. A similar phenomenon has been demonstrated before, and NMDA-dependent plasticity in the dentate gyrus and dorsal CA1 was shown to be the molecular mechanism beyond the observed effect [25, 26]. In the case of the “Modified” group, mice were forced to use flexible navigation from the onset of learning, since in order to get the reward they were following randomly changing contexts. Their attention was highly focused on extramaze cues, as there were sequence arrangements requiring consecutive entry into the same arm. It is possible that this high attention necessity may be the reason for stronger hippocampal engagement. It is worth noticing that in the “Modified” group, the activity of the rostral part of agranular retrosplenial cortex (RSA) is diminished, as compared to the “Alternation” group. This observation is in accordance with the previously reported negative feedback between the output fields of the hippocampal formation and the RSC. These direct inhibitory connections were first described on the anatomical level [7]. Functional studies confirmed these findings and revealed additional disynaptic inhibitory input [27–29]. According to these reports, the projections from CA1 and subiculum terminate in the granular retrosplenial area (RSG), but the region affected in our case is RSA. This effect might be due to the interactions between the two regions, as described in [30]. We also observed a gradient of activity along the rostro-caudal axis of the RSA. This confirms previously observed differential roles for rRSC and cRSC [24, 31], with the caudal part encoding the features of the environment, and the rostral area responsible for motor planning. The existence of a negative balance between HPC and RSC supports the concept that these structures may act competitively to drive the spatial behavior. Under the conditions where the hippocampus is engaged, it suppresses RSC in order to avoid conflicting behaviors. RSA is a likely candidate for harboring the alternative memory center. It receives sensory inputs from the visual cortex [32], and engages in an independent thalamocortical circuit [33]. The presence of molecular signature of a memory engram in this structure has been previously described by us and others [34, 35].

Notably, the RSC activity in the “Alternation” group is constantly high in all analyzed subregions, while the hippocampus is relatively inactive under these conditions. Animals in this group have access to exactly the same variety of external cues but during training it is displayed in sequence preserving natural tendency to alternate. A number of recent reports provide ample evidence that RSC could support spatial memory and navigation by itself. It encodes the animal’s position [36, 37], trajectory progression [38], head turning direction [39, 40], goal location [41] but also a number of unique spatial correlates [42], including motion- and direction independent heading signal [43] which suggests the existence of purely egocentric spatial code [44]. RSC could therefore shift rapidly between allocentric and egocentric spatial representations in order to flexibly support navigation. Indeed, lesion and inactivation data suggest the involvement of RSC in a number of spatial tasks, but its disruption rarely leads to a complete deficit [20, 34, 45–47]. Based on our current results, we postulate that a specific training protocol optimized for RSC engagement could minimize the involvement of the hippocampus and allow for easy dissecting of the alternate spatial memory circuits.

